# Impact of explicit area scaling on kinetic models involving multiple compartments

**DOI:** 10.1101/2020.10.30.361717

**Authors:** Pascal Holzheu, Ruth Großeholz, Ursula Kummer

## Abstract

**Background:** Computational modelling of cell biological processes is a frequently used technique to analyse the underlying mechanisms and to generally understand the behaviour of these processes in the context of a pathway, network or even the whole cell. The most common technique in this context is the usage of ordinary differential equations (ODEs) that describe the kinetics of the
relevant processes in mechanistic detail. Here, it is usually assumed that the content of the cell is well-stirred and thus homogeneous - which is of course an over-simplification, but often worked in the past. However, many processes happen at membranes and thus not in 3D, but in 2D. The scaling of the rates of these processes poses a special problem, if volumes of compartments are changed. They will typically scale with an area, but not with the volume of the involved compartment. However, commonly, this is neglected when setting up models and/or volume scaling also sometimes automatically happens when using modelling software in the field.

**Results:** Here, we investigate generic as well as specific, realistic cases to find out, how strong the impact of the wrong scaling is for the outcome of simulations. We show that the importance of correct area scaling depends on the architecture of the reaction site and its changes upon volume alterations and it is hard to foresee, if it has a significant impact or not just by looking at the original model set-up. Moreover, scaled rates might exhibit more or less control over the behaviour of the system and therefore, accordingly, incorrect scaling will have more or less influence.

**Conclusions:** Working with multi-compartment reactions requires a careful consideration of the correct scaling of the rates when changing the volumes of the involved compartments. The error following incorrect scaling - often done by scaling with the volume of the respective compartments can lead to significant aberrations of model behaviour.

## Background

The spatial separation of biological processes into different compartments constitutes one of the central features of eukaryotic cells. It allows for a specific control of pH, oxidative state and metabolite concentrations as well as keeping transcriptional and translational processes separated. Furthermore, ionic gradients across organellar membranes can be utilized for energy conversion as it is done by the proton gradient across the inner mitochondrial membrane, and nuclear signaling pathways require downstream factors to cross the nuclear membrane to reach their target. With compartmentalization playing such a central role in many processes, it comes as no surprise that researchers trying to understand these processes must take trans-membrane transport and other membrane bound processes into account. As most biological processes are complex to a degree that a pure qualitative description of what is happening in a specific situation is not sufficient to fully understand the system in question, computational modeling has become an indispensable part of biological research [1]. A variety of modeling approaches exists. A commonly used description of metabolic networks on a higher level of abstraction describing flux distributions has become popular in recent years - whole genome-scale metabolic models that are based on stoichiometric information and do not require kinetic detail [2]. However, using more coarse grained information also limits the analysis and predictions of such models to more general results. Other, less frequently used computational modeling approaches include stochastic formalism [3], partial differential equations (PDEs) [4], bayesian networks [5], boolean descriptions [6] and Petri-Nets [7] to name just a few. The most common formalism used in computational modeling however are ordinary differential equations (ODEs) [8]. These allow detailed mechanistic descriptions of the involved processes in a given system, but also require some existing knowledge of these mechanisms and the corresponding kinetic parameters or - alternatively - a lot of quantitative experimental data to determine at least some of the parameters via parameter fitting. While modeling received more and more attention, a need for a standardized format for storing and exchange of models in different modeling software arose. Therefore, the Systems Biology Markup Language (SBML) format was developed [9] which allows loading and analysis of computational models with different software and platforms. This format is especially suitable, but not restricted to ODEs.

ODEs express the time-dependent change of a substance concentration as a function of this concentration, the concentration of other involved substances in the system, as well as of kinetic parameters *p*:

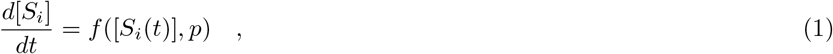

where *S_i_* is the vector of substance concentrations and *p* a vector of kinetic parameters. By means of numerical integration, the time-dependent concentrations of the respective substances can be obtained. During the process of constructing an ODE model, the modeler needs to carefully examine which molecular species and reactions to consider: On the one hand, all relevant processes of a system need to be captured. On the other hand, the model shouldn’t be unnecessarily complicated avoiding high computational costs during analysis and difficulties in being able to understand the behavior of the system in an adequate way. The same careful choices must be made when choosing the kinetic rate law for each reaction and information about the reaction mechanism is required during this process. In general, the velocity of a reaction is composed of the velocity of the forward reaction subtracted by the one of the backward reaction **expressed as concentration change**:

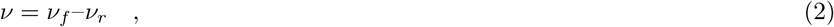

which in the simple case of a reversible mass-action-governed reaction such as

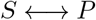

would read

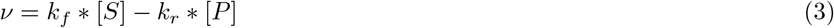

with equilibrium constant

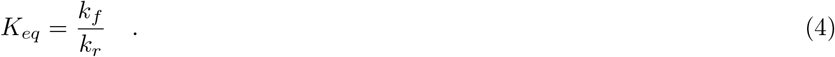

Here, [*S*] and [*P*] denote the substrate and product concentrations and *k_f_, k_r_* the kinetic constants of the forward and reverse reaction. More complicated kinetic terms, e.g. describing saturating rate laws as often caused by enzymatic catalysis like Michaelis-Menten or involving inhibitors and activators in the system are common in the cell biological context [10].

These expressions, like the above equation (3), are however only valid when the reactions occur within a single compartment. The reason for this is that the reaction velocity explicitly contains the concentrations of the reactants. If the reaction occurs across a membrane, such that *S* and *P* are present in different compartments with different volumes *V*_1_ and *V*_2_, respectively, the decrease of the concentration of *S* during a reaction instance does not lead to an equal increase of the concentration of *P*. To account for this, in many models, trans-membrane reactions are scaled with the volume ratio of the respective compartments (e.g. [11]). Another approach is required by SBML, where the usage of particle numbers instead of concentrations is the standard. Here, rates of transport reactions describe particle fluxes. This standard was selected to avoid problems as discussed below. However, in this case, again, one could be tempted to assume that for obtaining the reaction velocity **now in terms of changes in molar amounts**, it is sufficient to multiply the rates with the respective volumes of the compartments in which the respective substances reside in:

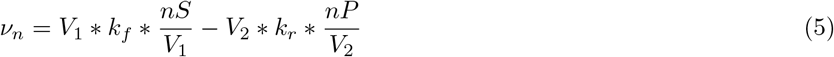

*nS, nP* being the molar amounts of *S* and *P*.

Indeed, both of the above scenarios (scaling with the volume ratio in the case of concentration changes or with the volumes in the case of molar amounts) can be found in publications and software in the field. However, in many, if not most cases the rate of a trans-membrane transport process will not scale with the volumes of the respective compartments - be it in the case of using particle numbers or in the case of concentrations. Instead, the transport rate scales with the amount of transporters present in the membrane, or in the case of diffusion of lipophilic substances through the membrane, simply with the area of the membrane. A very detailed description of different geometrical scenarios and the correct equations for each of these cases was recently published [12]. Therefore, the scaling of reaction rates should take into account the number of transporters in the membrane, or, assuming a constant transporter density, the surface area of the membrane. Thus, when changing volumes of compartments and subsequently adjusting the rate of a transport process, in the latter case, one has to compute the change in area A rather than taking into account the change in volume:

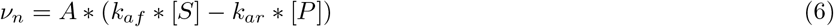

Depending on the exact nature of the architecture of the compartments and their interface the area might change at least in a non-linear correlation with the volume or it might not change at all, as we will discuss below. For example, when looking at spheroidal compartment such as the nucleus, the volume is approximately given as 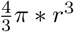, while the surface reads 4*π* * *r*^2^. Consequently, the difference in reaction velocities between the two scaling approaches in this case scales linearly with r. In other situations, the volume and surface of the compartment of interest can have a nonlinear relation, as a growth in surface does not necessarily go along with a similar change in volume.

The impact of the precise scaling on the system’s behaviour is of course also influenced by the sensitivity that the behaviour exhibits with respect to the exact reaction rate under consideration. Therefore, it can be useful to calculate the sensitivities of the model output regarding the rate of the transport reaction which is altered by the scaling. If there is only very little sencitivity and therefore large robustness with respect to this specific rate, the difference made by the exact scaling will also not be of major importance.

While a first-order reaction in a homogeneous system has the unit *s*^−1^, the need for area scaling can lead to unusual units of the resulting phenomenological kinetic parameters, as in the above described case: *m* * *s*^−1^ for *k_af_* and *k_ar_*. This is however no problem if one carefully assesses the meaning of the affected parameters.

Since what is stated above is not yet common practice among modelers, we investigate in the following the implications of not taking the surface area at which a transport reaction proceeds into account. We examined several published multicompartment models without area scaling. We further compared their behavior in diverse physiological circumstances, where volume ratios or the area to volume ratio change. Here, we demonstrate that the correct behavior can be obtained with any modeling approach as long as the parameters are treated accordingly.

## Methods

For model selection, we used the EBI BioModels database. From the curated models published there (last accessed: 10.07.2019), we selected the ones comprised of at least two different compartments with non-arbitrary volumes. Of those, we excluded models in which only the ratios between biological compartment volumes are considered as well as pharmacokinetic models.

The selected models were analysed and modified using the modeling software package COPASI, version 4.23 [13]. The time-courses of the relevant species’ concentrations were determined deterministically using LSODA as implemented in COPASI. Scaled sensitivities of steady-state concentrations and transient concentration maxima were calculated with COPASI as well.

## Results

### Vertical transport between root cells

Before studying the effects of different compartment conformations on models from the BioModels database, we first sought to understand the consequences of the different model structures in a simple toy model. Here, we used a model describing the vertical, reversible transport in *Arabidopsis thaliana* root epidermis cells. A particular characteristic of plant root cells is that they increase mostly in length and only to a small degree in width [14]: Young cells at the root tip (Fig. 1 - blue) are much smaller than mature cells in the older tissue (Fig. 1 - orange). Consequently, the vertical interaction surface between cells changes on a vastly different scale compared to either total cell surface or cell volume. Considering the fact that directional transport between cells is a crucial aspect to proper plant development [15], any model of such processes would have to account for the special characteristics of plant cells.

**Figure 1.**
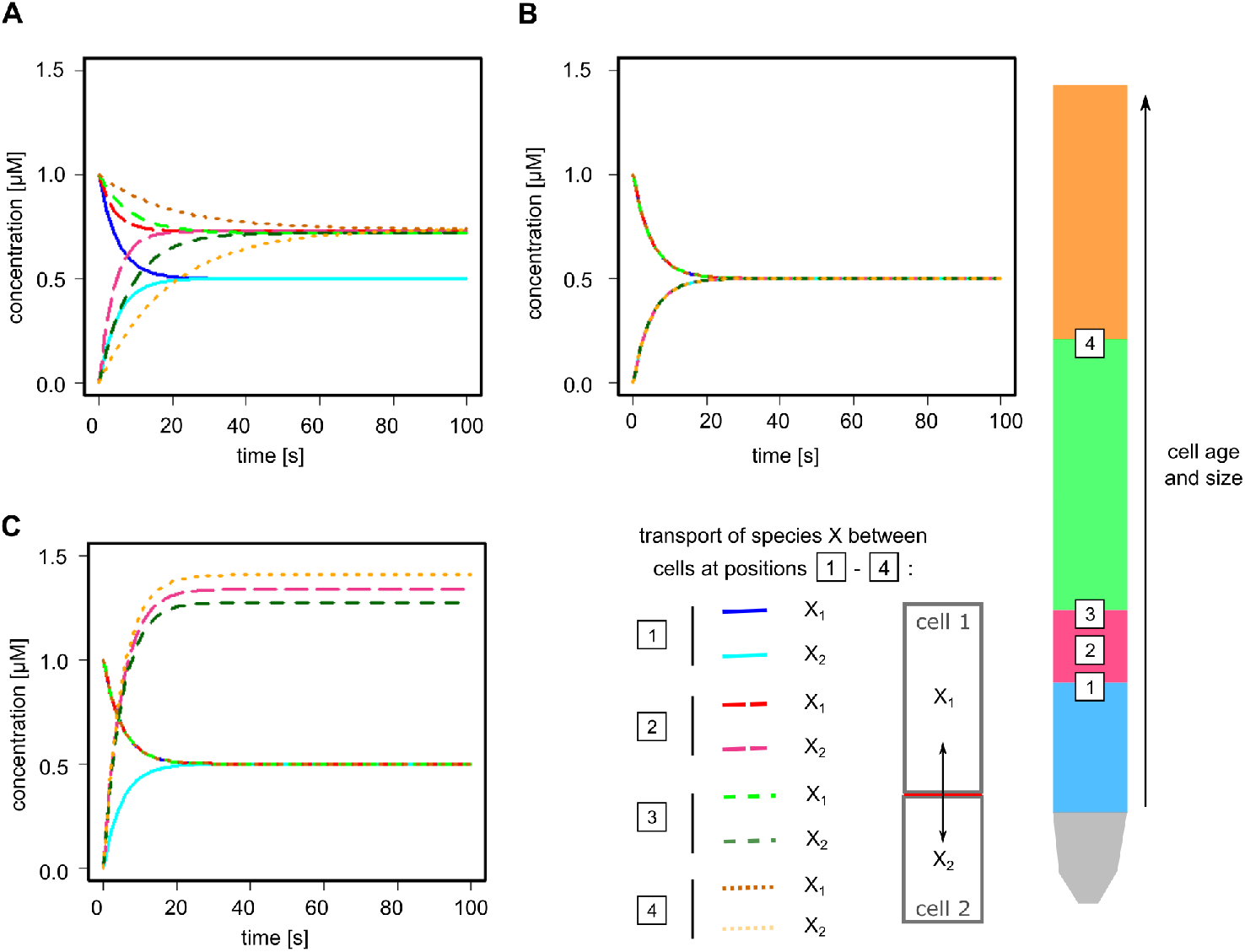
Impact of different modeling approaches on the vertical transport between root cells. The transport of species X was simulated for two epidermis cells at four different positions along the root vertical axis representing the different developmental zones: 1 - from the meristematic zone (blue, smallest cell volume and age) to the transition zone (red); 2 - within the transition zone (red); 3 - from the transition zone (red) to the elongation zone (green); 4 - from the elongation zone (green) to the maturation zone (orange, largest cell volume and age). A: Simulations of the area-scaling model. B: Simulations of the one compartment model. C: Simulations of the volume-scaling model. Parameters are listed in supporting table S1.

For this simplistic model, we analysed the transport of a species *X* between two epidermis cells in the *A. thaliana* root. This model comprises only one reaction:

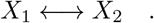

where *X*_1_ and *X*_2_ denote *X* in epidermis cells *A* and *B*, respectively. The cell sizes were chosen according to the measured dimensions of epidermis cells in the different root regions [14]. As we simulated the transport of *X*_1_ from an older and therefore larger cell to a younger (smaller) cell, we analysed four different transit situations (Fig. 1 - 1 to 4) that cover the whole range of cell sizes starting with the younger cells (transition 1 red → blue) to the oldest tissue (transition 4 orange → green).

We set the initial concentration of *X*_1_ to 1 *μ*M. This allows for an easy evaluation of not only the equilibrium concentrations of *X* in both cells but also of the speed, at which the equilibrium is reached. In particular, we analysed three different implementations of multi-compartment reactions: one universal compartment, multiple compartments with volume scaling and multiple compartments with area scaling using the membrane area between the cells. The parameters were adjusted so that the models exhibit identical behavior for scenario 1 (Fig. 1). For all subsequent simulations, the parameters were left unchanged to illustrate the differences of the different modelling assumptions.

As illustrated in Figure 1, the different implementations of the model react in different ways to changes in cell size. Here, the realistic area scaling model shows a clear delay in reaching the equilibrium as cell sizes increase (blue → red → green → orange) (Fig. 1A). Furthermore, the equilibrium concentrations of *X*_1_ and *X*_2_ shift on account of the different cell sizes. In contrast to this, the model consisting of a single compartment shows no change between the different scenarios (B). Last, the volume scaling model demonstrates a great shift in equilibrium concentrations as the transport rates are scaled with the individual compartment sizes (C). Unlike the area scaling model, *X* reaches the equilibrium at approximately the same time in the different scenarios. Altogether, this simple transport model already demonstrates that the consideration of the proper and adequate scaling can make a difference.

### Introducing Area Scaling to More Complex Models

Next, we analysed two realistic examples from the BioModels database. Here, we considered a RanGTP shuttling model [16], (BioModelsID 192) and a TGF-*β* model [17], (BioModelsID 342). To select these models we analysed the curated model entries of the BioModels database according to the compartmentalization. Here, we considered the number of compartments as well as the compartment size. Notably, only a small fraction of entries consisted of multi-compartment models with realistic compartment sizes (22 out of 825 curated models, see Table S1). The vast majority of models comprises only one compartment.

### Nuclear transport of HeLa cells is robust during growth

The RanGTP model by Görlich et al. (Fig. 2A) describes the shuttling of RanGTP from the nucleus to the cytoplasm and the transfer of RanGDP from the cytoplasm into the nucleus, the conversion reactions between RanGTP and RanGDP and the(un-)binding of nucleo-cytoplasmic shuttling factors (Fig. 2A) [16]. This model was used to demonstrate that the maintenance of a large RanGTP gradient requires a sufficiently large cytoplasm showing that compartment sizes themselves already have an impact on cellular behavior [16]. As the RanGTP concentration in the model is sensitive to changes in the transport reactions (see Table S3), we included a scaling factor describing the area of the nuclear envelope to scale these reactions with the nuclear area instead of the volume. In particular, there are two model reactions occurring between different compartments - the transport of RanGTP and RanGDP - are scaled with the nuclear volume in the original model. Here, we changed the kinetic rate laws of those two processes to a nuclear area scaling version, with the kinetic parameters *k_perm RanGDP_* and *k_perm RanGTP_* adjusted such as to reproduce the original model’s behavior (Fig. 2B).

**Figure 2.**
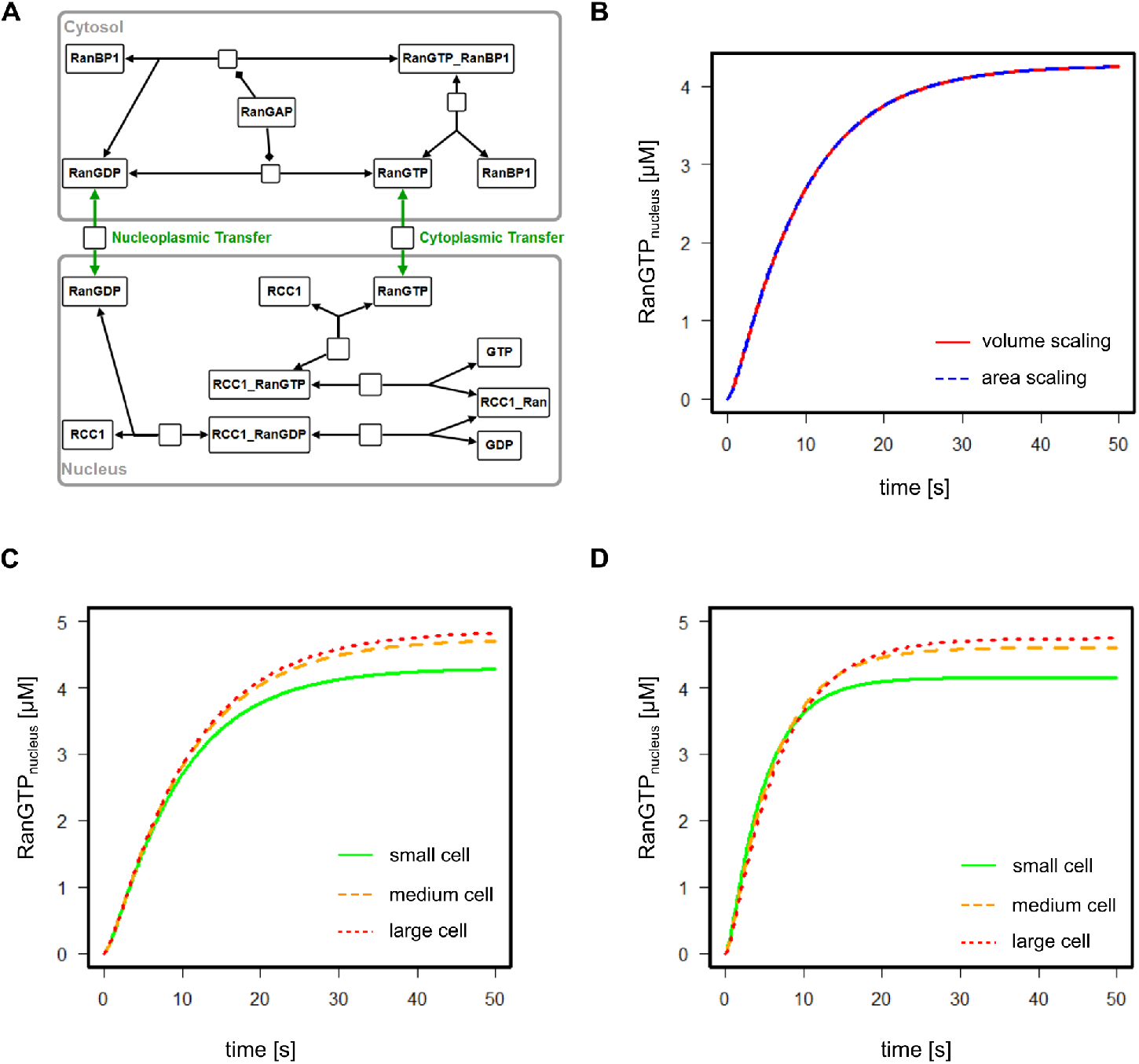
Analysis of the RanGTP shuttling model by Görlich et al.. A: Model reaction scheme according to the SBGN standard. B: Reproducing the behavior of the original model (red) with the area scaling model (blue). C: Simulations of the volume scaling model at different cell sizes. D: Simulations of the area scaling model at different cell sizes. Color code: green - small cell; orange - medium cell; red - large cell.

There are two model reactions occurring between different compartments - the transport of RanGTP and of RanGDP. These are scaled with the nuclear volume in the original model. Here, we changed the kinetic rate laws of those two processes to a nuclear area scaling version, with the kinetic parameters *k_perm RanGDP_* and *k_perm RanGTP_* adjusted such as to reproduce the original model’s behavior (Fig. 2B).

HeLa cells, upon which the model is based, undergo growth in both cytoplasm and nuclear volume during cell cycle progression. The cell volume varies on average between 2.2 and 5.2 * 10^3^ *μ*m^3^, whereas the nuclear volume varies between 1.6 and 2.6 * 10^3^ *μ*m^3^ [18]. Assuming that a larger cell volume corresponds to a larger nuclear volume (due to DNA duplication before cell division), this results in the compartment sizes listed in Table 1. We subsequently simulated the behavior of both the area scaling and the volume scaling model for all cell sizes listed in Table 1. Here, the volume scaling model (C) shows a slower dynamics compared to the area scaling model (D), both reaching the same steady state concentrations for the different cell sizes. The differences in these cases are not particularly significant, especially for the medium (red) and large (orange) cell sizes. This demonstrates that including the area as a scaling factor is not always necessary, especially if the qualitative outcome is more important than the quantitative. However, even though relative size changes of the two compartments in this example are small, there is already a notable difference which also points out that larger differences would result in significant differences in model behavior if unrealistic scaling (to the volume) is taken into account.

**Table 1.** Cytosolic and nuclear compartment sizes of the HeLa cells. Nuclear volume and surface area were calculated based on the nuclear cross-section [18].

|  |  | original model | small cell | medium cell | large cell |
| --- | --- | --- | --- | --- | --- |
| cell volume | $[\mu\text{m}^3]$ | $1.8 * 10^4$ | $2.2 * 10^3$ | $3.7 * 10^3$ | $5.2 * 10^3$ |
| nuclear cross section | $[\mu\text{m}^2]$ | NA | 155 | 200 | 245 |
| nuclear volume | $[\mu\text{m}^3]$ | $1.2 * 10^4$ | $1.45 * 10^3$ | $2.13 * 10^3$ | $2.88 * 10^3$ |
| nuclear surface | $[\mu\text{m}^2]$ | 2533.88 | 619.28 | 800.23 | 979.79 |

### Altered nuclear morphology in cancer cells affects signalling response

The model of Zi et al. describes the TGF-*β* induced Smad2 signaling pathway (Fig. 3A). The authors constructed the model to investigate the differential effects of variable TGF-*β*-doses on the intracellular signal dynamics, finding distinct responses of the cell to both sustained and pulsating TGF-*β*-stimulation.

**Figure 3.**
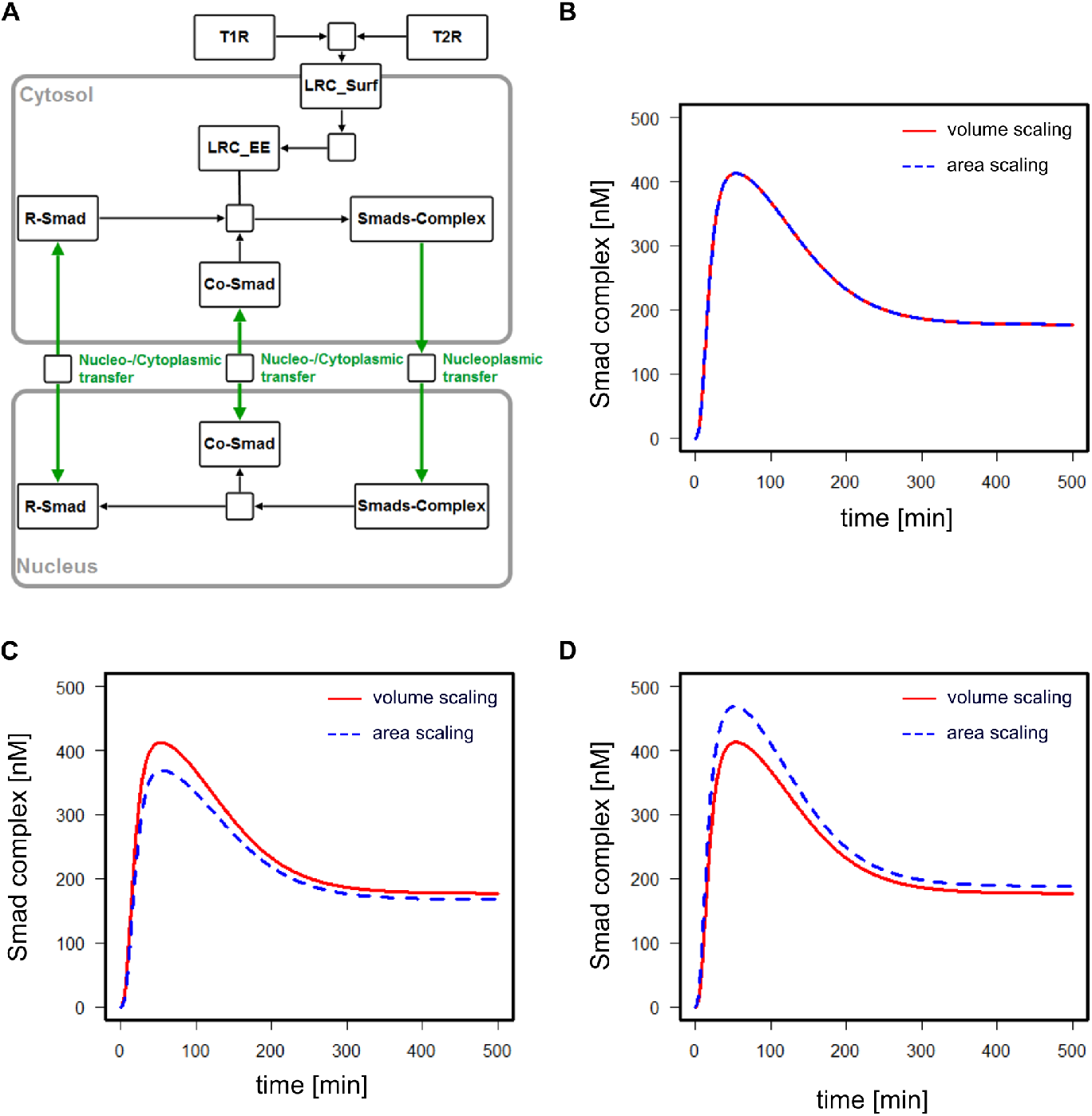
Analysis of the TGF-*β* induced Smad2 signaling pathway by Zi et al.. A: Model reaction scheme according to the SBGN standard. B: Reproducing the behavior of the original model (red) with the area scaling model (blue). C: Comparison of the simulations of the volume scaling model and the area scaling model at both increased cell and nuclear size. D: Comparison of the simulations of the volume scaling model and the area scaling model at increased nuclear surface area.

The model reactions include the binding and unbinding of TGF-*β* to T1R and T2R and their recycling at the plasma membrane, complex formation of R-Smad with Co-Smad and the shuttling of R-Smad, Co-Smad and the Smads-complex across the nuclear envelope. Similar to the RanGTP model, the processes of nuclear shuttling were changed from being scaled with the nuclear or cytoplamic volume to being scaled with the nuclear membrane area. The affected parameters were again adjusted to deliver the same output as before. The model was originally constructed using data from HaCaT cells, a human keratinocyte cell line. Skin cancer cells, as many other forms of cancer cells, often exhibit nuclei with irregular shapes as well as being bi- or multinucleated in several cases [19], both facts contributing to a higher surface area to volume ratio, thus posing a potential situation in which the differential behavior of volume- and area-scaled models can be observed.

The possible range of alterations here is large, so for the sake of this study, we analysed the effect of an increase of both the nuclear and cell radius by 50% (Fig. 3C) and the effect of a 50% increase of the nuclear surface area while keeping all volumes constant (Fig. 3D). This change affects only the area scaled model as in the volume-scaled version, the nuclear surface is not a parameter considered. Analogously to the example above, the parameters of the area-scaled model were adjusted to replicate the models original behavior (Fig. 3B) and we compared the simulations of both the original, volume-scaled model with the area-scaled one (Fig. 3C and 3D). Furthermore, the scaled sensitivities of the Smads-complex concentration to the transport reactions were calculated and shown to be sufficiently high (Table S4), suggesting that a change of the transport rates has a noticeable influence on the signaling output. It can be seen that both the steady state value of nuclear Smads-complex and especially its transient are different in both model versions, e.g. the peak concentration of nuclear Smads-complex in the area-scaled model is around 10% lower than in the volume-scaled version when both the cellular and nuclear volumes are increased by 50%, while its peak concentration is around 14% higher when just the nuclear area increased by 50%. Again, the differences are not huge. However, if the models serve a quantitative purpose they are significant enough.

This result corroborates that depending on the system under investigation, a careful consideration of the scaling of trans-compartmental rate laws can change the model predictions.

## Discussion

Kinetic modelling of biological systems has the potential to enhance our understanding of the respective processes. A good model can prevent unnecessary experiments and aid in the analysis of a system in states that are difficult to access experimentally. Computational models always represent a simplification of reality. Therefore, it is important to carefully examine the assumptions applied in the respective model as to not undermine its predictive power. These assumptions can concern the formulation of biochemical reactions, compartment sizes or the timescale of different processes. In particular, biochemical reaction velocities typically scale with the amount of enzymes present in the cell. For transport processes the rate of the reaction scales with the number of transporters. If no accurate measurements of transporter concentrations are available, the membrane area of the respective compartment can serve as proxy.

In this study, we examined the effects of changing cell sizes both through growth and other, e.g. carcinogenic processes on multi-compartment processes in ODE models. So far, the effects of cell size and shape have been examined in PDE models, where they influence the response in signaling pathways [20] and, on the macroscopic level, determine physiological as well as developmental outputs [21]. In contrast to this, the effects of changing cell shape or size on the simulations of ODE models have been largely ignored. While a great number of metabolic processes involve more than one compartment, the existence of different compartments is not always considered explicitly in the respective ODE model. To avoid the complications of including several compartments, pathways are often simplified to comprise only one compartment or omit the existence of the membrane.

In fact, the majority of curated models in the BioModels database consists of one compartment. This simplification implies by no means that the resulting model is wrong or in any way unable to describe the experimental data correctly. However, as the respective model comprises only one compartment or the membrane is not explicitly included as scaling factor, the kinetic parameters of any multi-compartment reaction will implicitly include that information. Therefore, these models are only correct for that particular cell size with that particular spatial conformation. Any change in the compartmental situation, i.e. cell shape, can necessitate the adaptation of model parameters - and not just the respective volumes themselves.

Depending on the modeled system, the effects of changing cell shape or size can affect the model behavior greatly. As illustrated by the model describing the vertical transport between *A. thaliana* root cells, growth can have a big impact on the speed of these processes. While this model nicely demonstrates the necessity of adjusting parameter values, not all cells grow as much as *A. thaliana* epidermis root cells that expand from an initial length of 8 *μ*m to up to 220 *μ*m in length. In fact, for the measured cell and nucleus sizes for HeLa cells [18], the behavior of the RanGTP shuttling model varies little between the different settings: While the RanGTP concentration is sensitive to changes in the transport reactions, the model response is quite robust as we change both reactions, nucleoplasmic and cytoplasmic transfer, concomitantly by the same factor.

In contrast to this, changing the compartment sizes caused a notable difference in the output of the TGF-*β* signaling model by Zi *et al*.. Not only is the Smad-complex concentration sufficiently sensitive to the transport reactions, but the model architecture also means that the change compartment size has a notable impact on the output function. In more general terms, this means that - depending on how the system is decoded (signal amplitude *versus* signal duration) - the area scaling model would transmit different information than the volume scaling model unless the parameters are carefully adjusted.

## Conclusions

Altogether, our analysis demonstrates that including the membrane as scaling factor - or at least carefully adjusting the parameters of multi-compartment reactions - can be necessary to observe the correct model behavior. Nonetheless, the exact impact of not adjusting the model on the simulated behavior depends on the modeled system itself, its geometry and the control that a transport reactions holds over the behavior of the system.

It is important to note that there are software tools that always automatically scale with the volume, if this is changed. According to the above said this is clearly wrong. Software like COPASI does correctly assume the user to know about particle fluxes and adjust these according to the impact that a changing volume of a compartment has. It also recently allowed for the definition of two-dimensional compartments, which should lead to more models considering the membrane as compartment or scaling-factor.

## Abbreviations

ODE: ordinary differential equations
PDE: partial differential equations
Ran: RAs-related Nuclear protein
SBML: Systems Biology Markup Language
TGF-*β*: Transforming Growth Factor - *β*

## Ethics approval and consent to participate

Not applicable

## Consent to publish

Not applicable

## Availability of data and materials

All computational models used in this study are available in the supporting material.

## Author’s contributions

PH, RG and UK conceived the idea for the study, PH and RG performed all computations and PH, RG and UK wrote the paper. All authors have read and approved the manuscript.

## Acknowledgements

Sven Sahle and Frank Bergmann are acknowledged for fruitful discussions. Particular thanks to Jannie Hofmeyr who pointed out the importance of area scaling years ago.

## Additional Files

### Supporting Information

List of curated, multi-compartment models in the biomodels database with realistic compartment sizes (Table S1). Detailed description of the models used in this study, including the setup and parameters of the transport model (Tables S2, S3), the changes to the RanGTP transport and the TGF-beta signaling model as well asthe calculated scaled sensitivity values for both models (Tables S4, S5).

### Model Files

Copasi files for all models used in this study.

## References

1. Noble, D.: The rise of computational biology. Nature Reviews Molecular Cell Biology 3(6), 459 (2002)

2. O’Brien, E.J., Monk, J.M., Palsson, B.O.: Using genome-scale models to predict biological capabilities. Cell 161(5), 971–987 (2015)

3. Wilkinson, D.J.: Stochastic modelling for systems biology. Chapman and Hall/CRC (2006)

4. Eungdamrong, N.J., Iyengar, R.: Modeling cell signaling networks. Biology of the Cell 96(5), 355–362 (2004)

5. Sachs, K., Gifford, D., Jaakkola, T., Sorger, P., Lauffenburger, D.A.: Bayesian network approach to cell signaling pathway modeling. Sci. STKE 2002(148), 38–38 (2002)

6. Saadatpour, A., Albert, R.: Boolean modeling of biological regulatory networks: a methodology tutorial. Methods 62(1), 3–12 (2013)

7. Chaouiya, C.: Petri net modelling of biological networks. Briefings in bioinformatics 8(4), 210–219 (2007)

8. Hübner, K., Sahle, S., Kummer, U.: Applications and trends in systems biology in biochemistry. The FEBS journal 278(16), 2767–2857 (2011)

9. Hucka, M., Finney, A., Sauro, H.M., Bolouri, H., Doyle, J.C., Kitano, H., Arkin, A.P., Bornstein, B.J., Bray, D., Cornish-Bowden, A., et al.: The systems biology markup language (sbml): a medium for representation and exchange of biochemical network models. Bioinformatics 19(4), 524–531 (2003)

10. Liebermeister, W., Klipp, E.: Bringing metabolic networks to life: Convenience rate law and thermodynamic constraints. Theoretical Biology and Medical Modelling 3, 41 (2006). doi:10.1186/1742-4682-3-41

11. Kholodenko, B.N., Demin, O.V., Moehren, G., Hoek, J.B.: Quantification of short term signaling by the epidermal growth factor receptor. Journal of Biological Chemistry 274(42), 30169–30181 (1999)

12. Hofmeyr, J.-H.S.: Kinetic modelling of compartmentalised reaction networks. BioSystems 197, 104203 (2020)

13. Hoops, S., Sahle, S., Gauges, R., Lee, C., Pahle, J., Simus, N., Singhal, M., Xu, L., Mendes, P., Kummer, U.: Copasi—a complex pathway simulator. Bioinformatics 22(24), 3067–3074 (2006)

14. van Esse, G.W., Westphal, A.H., Surendran, R.P., Albrecht, C., van Veen, B., Borst, J.W., de Vries, S.C.: Quantification of the brassinosteroid insensitive1 receptor in planta. Plant physiology 156(4), 1691–1700 (2011)

15. Friml, J.: Auxin transport—shaping the plant. Current opinion in plant biology 6(1), 7–12 (2003)

16. Görlich, D., Seewald, M.J., Ribbeck, K.: Characterization of Ran-driven cargo transport and the RanGTPase system by kinetic measurements and computer simulation. The EMBO journal 22(5), 1088–100 (2003). doi:10.1093/emboj/cdg113

17. Zi, Z., Feng, Z., Chapnick, D.A., Dahl, M., Deng, D., Klipp, E., Moustakas, A., Liu, X.: Quantitative analysis of transient and sustained transforming growth factor-*β* signaling dynamics. Molecular systems biology 7(1), 492 (2011). doi:10.1038/msb.2011.22

18. Puck, T.T., Marcus, P.I., Cieciura, S.J.: Clonal growth of mammalian cells in vitro. Journal of Experimental Medicine 103(2), 273–284 (1956). doi:10.1084/jem.103.2.273. http://jem.rupress.org/content/103/2/273.full.pdf

19. Dey, P.: Cancer nucleus: morphology and beyond. Diagnostic cytopathology 38(5), 382–390 (2010)

20. Meyers, J., Craig, J., Odde, D.J.: Potential for control of signaling pathways via cell size and shape. Current biology 16(17), 1685–1693 (2006)

21. Baker, R.E., Gaffney, E., Maini, P.: Partial differential equations for self-organization in cellular and developmental biology. Nonlinearity 21(11), 251 (2008)

