## Supporting Information for "Impact of explicit area scaling on kinetic models involving multiple compartments"

### 1 Models with realistic compartment sizes from the BioModels databse

**Table 1 Curated multi-compartment models in the biomodels database. Listed are only those models with realistic compartment sizes with biomodel number (#) [1] (https://wwwdev.ebi.ac.uk/biomodels/, last accessed: 14.11.2019). The chosen example models are highlighted in bold font. Models built in Virtual Cell [2] are marked by VC.**

| # | Description | Reference |
|---|---|---|
| 16 | Circadian oscillations in the *drosophila* period protein (PER) | [3] |
| 19 | Dynamics of the MAP kinase cascade activated by surface and internalized EGF receptors | [4] |
| 59 | $Ca^{2+}$ flux in pancreatic beta-cells: role of the plasma membrane and intracellular stores | [5] |
| 75 (VC) | Kinetic analysis of receptor-activated phosphoinositide turnover | [6] |
| 122, 123 | NFAT and NF$\kappa$B activation in T lymphocytes | [7] |
| 161 (VC) | Compartment-specific feedback loop and regulated trafficking can result in sustained activation of Ras at the Golgi | [8] |
| 162 (VC) | Modeling and analysis of calcium signaling events leading to long-term depression in cerebellar Purkinje cells. | [9] |
| 173 | Smad nucleocytoplasmic shuttling as a dynamic signal-interpreting system | [10] |
| 182 (VC) | Cell shape and negative links in regulatory motifs together control spatial information flow in signaling networks. | [11] |
| **192** | Ran-driven cargo transport and the RanGTPase system | [12] |
| 250 | Ligand-specific c-Fos expression | [13] |
| **342** | Transient and sustained transforming growth factor-$\beta$ signaling | [14] |
| 399 | Thermodynamically consistent model calibration in chemical kinetics | [15] |
| 407 | Heterogeneity reduces sensitivity of cell death for TNF-stimuli | [16] |
| 474 | Regulation of insulin signaling by oxidative stress | [17] |
| 482 | The selective control of glycolysis, gluconeogenesis and glycogenesis by temporal insulin patterns | [18] |
| 505, 506 | Fatty acid $\beta$-oxidation | [19] |
| 547 | Arsenic transport, distribution and detoxification in yeast | [20] |
| 610 | Quantitavie analysis of glycerol accumulation, glycolysis and growth under hyper osmotic stress | [21] |
| 638 | Yeast GPCR signaling reflects the fraction of occupied receptors | [22] |
| 699, 702 | Regulation of Tem1 by the GAP complex in Spindle Position Checkpoint | [23] |
| 705, 706 | Response of FOXO transcription factors to multiple post-translational modifications made by ageing-related signaling pathways | [24] |

#### 2 Setup of the transport model

The model comprises only one reversible reaction:

$X_1 \leftrightarrow X_2$

In the following equations, concentrations are noted as $[X]$, $V$ refers to cellular volumes, $A_{interaction}$ defines the interaction area between cells and $k_{1,2,A1,A2}$ are kinetic parameters.

*One compartment*

$$-\frac{[X_1] \cdot V_{cell}}{dt} = \frac{[X_2] \cdot V_{cell}}{dt} = V_{cell} * (k_1 * [X_1] - k_2 * [X_2]) \quad (1)$$

*Volume scaling transport*

$$-\frac{[X_1] \cdot V_{cell\,1}}{dt} = \frac{[X_2] \cdot V_{cell\,2}}{dt} = V_{cell\,1} * k_1 * [X_1] - V_{cell\,2} * k_2 * [X_2] \quad (2)$$

*Area scaling transport*

$$-\frac{[X_1] \cdot V_{cell\,1}}{dt} = \frac{[X_2] \cdot V_{cell\,2}}{dt} = A_{interaction} * (k_{A1} * [X_1] - k_{A2} * [X_2]) \quad (3)$$

#### 3 Parameters of the transport model

**Table 2 Parameters of the transport model for the first transition (transition to meristematic zone) and all three modeling approaches.**

| model structure | model entity | value |
|---|---|---|
| one compartment | $k_1$ | $0.1\,s^{-1}$ |
| | $k_2$ | $0.1\,s^{-1}$ |
| volume scaling | $k_1$ | $0.1\,s^{-1}$ |
| | $k_2$ | $0.1\,s^{-1}$ |
| area scaling | $k_{A1}$ | $8.8192 * 10^{-6}\,dm * s^{-1}$ |
| | $k_{A2}$ | $8.8192 * 10^{-6}\,dm * s^{-1}$ |

**Table 3 Compartment sizes of the different transitions for the toy model. For the one compartment model, the cell has the volume listed in the column cell 1. The interaction area was only used for the area-scaling model.**

| transition | cell 1 | cell 2 | interaction area |
|---|---|---|---|
| 1 | $8.47 * 10^{-13}\,l$ | $8.47 * 10^{-13}\,l$ | $9.604 * 10^{-9}\,dm^2$ |
| 2 | $2.271 * 10^{-12}\,l$ | $8.47 * 10^{-13}\,l$ | $1.69 * 10^{-8}\,dm^2$ |
| 3 | $5.803 * 10^{-12}\,l$ | $2.271 * 10^{-12}\,l$ | $2.1904 * 10^{-8}\,dm^2$ |
| 4 | $1.6364 * 10^{-11}\,l$ | $5.803 * 10^{-12}\,l$ | $2.5281 * 10^{-8}\,dm^2$ |

#### 4 RanGTP transport model

##### 4.1 Altered reactions

*Cytoplasmic transfer* $(RanGTP_{nuc} \leftrightarrow RanGTP_{cy})$

new reaction rate law:

$k_{A\,perm\,RanGTP} * A_{nucleus} * (RanGTP_{nuc} - RanGTP_{cy})$

with $k_{A\,perm\,RanGTP}$: $1.429 * 10^{-6}\,dm * s^{-1}$

$Nucleoplasmic\ transfer \quad (RanGDP_{nuc} \leftrightarrow RanGDP_{cy})$
new reaction rate law:
$k_{A\,perm\,RanGDP} * A_{nucleus} * (RanGDP_{nuc} - RanGDP_{cy})$
with $k_{A\,perm\,RanGDP}$: $5.714 * 10^{-6}\,dm * s^{-1}$

##### 4.2 Sensitivities

**Table 4 Scaled sensitivities of the nuclear RanGTP concentration with respect to model parameters at steady state in the original, volume-scaling model by Görlich et al..**

| reaction | parameter | scaled sensitivity |
|---|---|---|
| RCC1_binding | $k_1$ | $0.139263$ |
| RCC1_binding | $k_2$ | $-0.13091$ |
| GDP_dissociation | $k_1$ | $0.133022$ |
| GDP_dissociation | $k_2$ | $-0.110622$ |
| GTP_binding | $k_1$ | $0.112722$ |
| GTP_binding | $k_2$ | $-0.111463$ |
| RanGTP_release | $k_1$ | $0.145492$ |
| RanGTP_release | $k_2$ | $-0.144969$ |
| Cytoplasmic_transfer | $k_{A\,perm\,RanGTP}$ | $-0.257474$ |
| Nucleoplasmic_transfer | $k_{A\,perm\,RanGDP}$ | $0.223293$ |
| RanGTP_binding | $k_1$ | $5.06681 * 10^{-05}$ |
| RanGTP_binding | $k_2$ | $-2.68329 * 10^{-10}$ |
| RanBP1_RanGDP | $K_m$ | $-1.2809 * 10^{-05}$ |
| RanBP1_RanGDP | $k_{cat}$ | $1.2804 * 10^{-05}$ |
| RanGAP_RanGDP | $K_{m\,GAP}$ | $-0.00109736$ |
| RanGAP_RanGDP | $k_{cat\,GAP}$ | $0.00110842$ |

#### 5 Smad2 signalling model

##### 5.1 altered reactions

$R1\_Smad2\_import \quad (Smad2\_c \rightarrow Smad2\_n)$
new reaction rate law:
$A_{nucleus} * k_{A\,imp}\_Smad2c * Smad2c$
with $k_{A\,imp}\_Smad2c$: $7 * 10^{-3}\,dm * s^{-1}$

$R2\_Smad2\_export \quad (Smad2\_n \rightarrow Smad2\_c)$
new reaction rate law:
$A_{nucleus} * k_{A\,exp}\_Smad2n * Smad2n$
with $k_{A\,exp}\_Smad2n$: $1.4583 * 10^{-2}\,dm * s^{-1}$

$R3\_Smad4\_import \quad (Smad4\_c \rightarrow Smad4\_n)$
new reaction rate law:
$A_{nucleus} * k_{A\,imp}\_Smad4c * Smad4c$
with $k_{A\,imp}\_Smad4c$: $3.5 * 10^{-3}\,dm * s^{-1}$

$R4\_Smad4\_export \quad (Smad4\_n \rightarrow Smad4\_c)$
new reaction rate law:
$A_{nucleus} * k_{A\,exp}\_Smad4c * Smad4c$
with $k_{A\,exp}\_Smad4c$: $7.2917 * 10^{-3}\,dm * s^{-1}$

$R24\_Smads\_Complex\_import \quad (Smads\_Complex\_c \rightarrow Smads\_Complex\_n)$
new reaction rate law:
$A_{nucleus} * k_{A\,imp}\_Smads\_Complex\_c * Smads\_Complex\_c$
with $k_{A\,imp}\_Smads\_Complex\_c$: $7 * 10^{-3}\,dm * s^{-1}$

#### 5.2 Sensitivities

**Table 5 Scaled sensitivities of the maximal nuclear Smads_Complex concentration with respect to model parameters in the original, volume-scaling model.**

| reaction | parameter | scaled sensitivity |
|---|---|---|
| R1_Smad2_import | $k_{A\,imp}\_Smad2c$ | $-0.0666527$ |
| R2_Smad2_export | $k_{A\,exp}\_Smad2n$ | $0.0916061$ |
| R3_Smad4_import | $k_{A\,imp}\_Smad4c$ | $-0.0666877$ |
| R4_Smad4_export | $k_{A\,exp}\_Smad4n$ | $0.0819686$ |
| R5_T1R_production | $v_{T1R}$ | $0.024528$ |
| R6/R12/R18_T1R/T2R/LRC_Cave_formation | $ki_{Cave}$ | $-0.340296$ |
| R7/R13/R19_T1R/T2R/LRC_Cave_recycling | $kr_{Cave}$ | $0.0842739$ |
| R8/R14/R20_T1R/T2R/LRC_EE_formation | $ki_{EE}$ | $0.302112$ |
| R9/R15/R21_T1R/T2R/LRC_EE_recycling | $kr_{EE}$ | $-0.130795$ |
| R10_T1R_EE_degradation | $k_{deg}\_T1R\_EE$ | $-0.00708498$ |
| R11_T2R_production | $v_{T2R}$ | $0.129776$ |
| R16_T2R_EE_degradation | $k_{deg}\_T2R\_EE$ | $-0.0224511$ |
| R17_LRC_formation | $k_{LRC}$ | $0.0242155$ |
| R22_LRC_EE_degradation | $k_{cd}$ | $-0.0550185$ |
| R23_Smads_Complex_formation | $k\_Smads\_Complex\_c$ | $0.486002$ |
| R24_Smads_Complex_import | $k_{A\,imp}\_Smads\_Complex_c$ | $0.198014$ |
| R25_Smads_Complex_Dissociation | $k_{diss}\_Smads\_Complex\_n$ | $-0.722791$ |
| R26_LRC_Cave_degradation | $k_{lid}$ | $-0.00316132$ |

**Author details**

**References**


1. Li, C., Donizelli, M., Rodriguez, N., Dharuri, H., Endler, L., Chelliah, V., Li, L., He, E., Henry, A., Stefan, M.I., Snoep, J.L., Hucka, M., Le Novère, N., Laibe, C.: BioModels Database: An enhanced, curated and annotated resource for published quantitative kinetic models. BMC Systems Biology **4**(1), 92 (2010). doi:10.1186/1752-0509-4-92
2. Schaff, J., Fink, C.C., Slepchenko, B., Carson, J.H., Loew, L.M.: A general computational framework for modeling cellular structure and function. Biophysical Journal **73**(3), 1135–1146 (1997). doi:10.1016/S0006-3495(97)78146-3
3. Goldbeter, A.: A model for circadian oscillations in the Drosophila period protein (PER). Proceedings. Biological sciences **261**(1362), 319–24 (1995). doi:10.1098/rspb.1995.0153
4. Schoeberl, B., Eichler-Jonsson, C., Gilles, E.D., Müller, G.: Computational modeling of the dynamics of the MAP kinase cascade activated by surface and internalized EGF receptors. Nature biotechnology **20**(4), 370–5 (2002). doi:10.1038/nbt0402-370
5. Fridlyand, L.E., Tamarina, N., Philipson, L.H.: Modeling of Ca2+ flux in pancreatic beta-cells: role of the plasma membrane and intracellular stores. American journal of physiology. Endocrinology and metabolism **285**(1), 138–54 (2003). doi:10.1152/ajpendo.00194.2002

6. Xu, C., Watras, J., Loew, L.M.: Kinetic analysis of receptor-activated phosphoinositide turnover. The Journal of cell biology **161**(4), 779–91 (2003). doi:10.1083/jcb.200301070
7. Fisher, W.G., Yang, P.-C., Medikonduri, R.K., Jafri, M.S.: NFAT and NFkappaB activation in T lymphocytes: a model of differential activation of gene expression. Annals of biomedical engineering **34**(11), 1712–28 (2006). doi:10.1007/s10439-006-9179-4
8. Eungdamrong, N.J., Iyengar, R.: Compartment-specific feedback loop and regulated trafficking can result in sustained activation of Ras at the Golgi. Biophysical journal **92**(3), 808–15 (2007). doi:10.1529/biophysj.106.093104
9. Hernjak, N., Slepchenko, B.M., Fernald, K., Fink, C.C., Fortin, D., Moraru, I.I., Watras, J., Loew, L.M.: Modeling and analysis of calcium signaling events leading to long-term depression in cerebellar Purkinje cells. Biophysical journal **89**(6), 3790–806 (2005). doi:10.1529/biophysj.105.065771
10. Schmierer, B., Tournier, A.L., Bates, P.A., Hill, C.S.: Mathematical modeling identifies Smad nucleocytoplasmic shuttling as a dynamic signal-interpreting system. Proceedings of the National Academy of Sciences of the United States of America **105**(18), 6608–13 (2008). doi:10.1073/pnas.0710134105
11. Neves, S.R., Tsokas, P., Sarkar, A., Grace, E.A., Rangamani, P., Taubenfeld, S.M., Alberini, C.M., Schaff, J.C., Blitzer, R.D., Moraru, I.I., Iyengar, R.: Cell shape and negative links in regulatory motifs together control spatial information flow in signaling networks. Cell **133**(4), 666–80 (2008). doi:10.1016/j.cell.2008.04.025
12. Görlich, D., Seewald, M.J., Ribbeck, K.: Characterization of Ran-driven cargo transport and the RanGTPase system by kinetic measurements and computer simulation. The EMBO journal **22**(5), 1088–100 (2003). doi:10.1093/emboj/cdg113
13. Nakakuki, T., Birtwistle, M.R., Saeki, Y., Yumoto, N., Ide, K., Nagashima, T., Brusch, L., Ogunnaike, B.A., Okada-Hatakeyama, M., Kholodenko, B.N.: Ligand-specific c-Fos expression emerges from the spatiotemporal control of ErbB network dynamics. Cell **141**(5), 884–96 (2010). doi:10.1016/j.cell.2010.03.054
14. Zi, Z., Feng, Z., Chapnick, D.A., Dahl, M., Deng, D., Klipp, E., Moustakas, A., Liu, X.: Quantitative analysis of transient and sustained transforming growth factor-$\beta$ signaling dynamics. Molecular systems biology **7**(1), 492 (2011). doi:10.1038/msb.2011.22
15. Jenkinson, G., Goutsias, J.: Thermodynamically consistent model calibration in chemical kinetics. BMC systems biology **5**(1), 64 (2011). doi:10.1186/1752-0509-5-64
16. Schliemann, M., Bullinger, E., Borchers, S., Allgöwer, F., Findeisen, R., Scheurich, P.: Heterogeneity reduces sensitivity of cell death for TNF-stimuli. BMC systems biology **5**(1), 204 (2011). doi:10.1186/1752-0509-5-204
17. Smith, G.R., Shanley, D.P.: Computational modelling of the regulation of Insulin signalling by oxidative stress. BMC systems biology **7**(1), 41 (2013). doi:10.1186/1752-0509-7-41
18. Noguchi, R., Kubota, H., Yugi, K., Toyoshima, Y., Komori, Y., Soga, T., Kuroda, S.: The selective control of glycolysis, gluconeogenesis and glycogenesis by temporal insulin patterns. Molecular Systems Biology **9**(1), 664 (2013). doi:10.1038/msb.2013.19
19. van Eunen, K., Simons, S.M.J., Gerding, A., Bleeker, A., den Besten, G., Touw, C.M.L., Houten, S.M., Groen, B.K., Krab, K., Reijngoud, D.-J., Bakker, B.M.: Biochemical competition makes fatty-acid $\beta$-oxidation vulnerable to substrate overload. PLoS computational biology **9**(8), 1003186 (2013). doi:10.1371/journal.pcbi.1003186
20. Talemi, S.R., Jacobson, T., Garla, V., Navarrete, C., Wagner, A., Tamás, M.J., Schaber, J.: Mathematical modelling of arsenic transport, distribution and detoxification processes in yeast. Molecular microbiology **92**(6), 1343–56 (2014). doi:10.1111/mmi.12631
21. Petelenz-Kurdziel, E., Kuehn, C., Nordlander, B., Klein, D., Hong, K.K., Jacobson, T., Dahl, P., Schaber, J., Nielsen, J., Hohmann, S., Klipp, E.: Quantitative Analysis of Glycerol Accumulation, Glycolysis and Growth under Hyper Osmotic Stress. PLoS Computational Biology **9**(6), 1003084 (2013). doi:10.1371/journal.pcbi.1003084
22. Bush, A., Vasen, G., Constantinou, A., Dunayevich, P., Patop, I.L., Blaustein, M., Colman-Lerner, A.: Yeast GPCR signaling reflects the fraction of occupied receptors, not the number. Molecular systems biology **12**(12), 898 (2016)
23. Caydasi, A.K., Lohel, M., Grünert, G., Dittrich, P., Pereira, G., Ibrahim, B.: A dynamical model of the spindle position checkpoint. Molecular systems biology **8**, 582 (2012). doi:10.1038/msb.2012.15
24. Smith, G.R., Shanley, D.P.: Modelling the Response of FOXO Transcription Factors to Multiple Post-Translational Modifications Made by Ageing-Related Signalling Pathways. PLoS ONE **5**(6), 11092 (2010). doi:10.1371/journal.pone.0011092
